# Defining melanoma combination therapies that provide senolytic sensitivity in human melanoma cells

**DOI:** 10.1101/2023.10.01.560354

**Authors:** Daméhan Tchelougou, Nicolas Malaquin, Guillaume Cardin, Jordan Desmul, Simon Turcotte, Francis Rodier

## Abstract

Malignant Melanoma that resists immunotherapy remains the deadliest form of skin cancer owing to poor clinically lasting responses. Alternative like genotoxic or targeted chemotherapy trigger various cancer cell fates after treatment including cell death and senescence. Senescent cells can be eliminated using senolytic drugs and we hypothesize that the targeted elimination of therapy-induced senescent melanoma cells could complement both conventional and immunotherapies.

We utilized a panel of cells representing diverse mutational background relevant to melanoma and found that they developed distinct senescence phenotypes in response to treatment. A genotoxic combination therapy of carboplatin-paclitaxel or irradiation triggered a mixed response of cell death and senescence, irrespective of BRAF mutation profiles. DNA damage-induced senescent cells exhibited morphological changes, residual DNA damage, and increased senescence-associated secretory phenotype (SASP). In contrast, dual targeted inhibition of Braf and Mek triggered a partially reversible senescence-like state without DNA damage or SASP.

To assess the sensitivity to senolytics we employed a novel real-time imaging-based death assay and observed that Bcl-xl/Bcl-2 inhibitors and piperlongumine were effective in promoting death of carboplatin-paclitaxel and irradiation-induced senescent melanoma cells, while senescent-like cells resulting from Braf-Mek inhibition remained unresponsive. Interestingly, a direct synergy between Bcl-2/Bcl-xl inhibitors and Braf-Mek inhibitors was observed when used out the context of senescence. Overall, we highlight that the hallmarks of melanoma senescence and sensitivity to senolytics are context dependent and provide evidence of effective combinations of senotherapy drugs that could reduce treatment resistance while also discussing the limitations of this strategy in human melanoma cells.

## 1 Introduction

Malignant melanoma remain the leading cause of skin cancer death worldwide including North America, Europe, and Australia, despite the revolutionary impact that immunotherapy has provided for its treatment (Ferlay et al., 2021, Bray et al., 2018, Siegel et al., 2020). Notably, hotspot mutations in the mitogen-activated protein kinase (MAPK) pathway, namely BRAF and NRAS, are frequently observed in approximately 50% and 25% of melanoma patients, respectively (Davies et al., 2002, Edlundh-Rose et al., 2006). Targeted therapies against these oncogenic mutations, such as small molecule inhibitors of MAPK kinase kinases (MEK) and BRAF, have shown significant benefits; however, the majority of patients eventually develop drug resistance and experience disease relapse (Perez-Lorenzo and Zheng, 2012).

Melanoma often harbors additional mutations in tumor suppressor genes such as TP53 (p53) and CDKN2A (p16) (Hodis et al., 2012, Shain and Bastian, 2016), which play crucial roles in regulating cellular senescence. Although recent therapies targeting immune checkpoint proteins have exhibited remarkable clinical efficacy, only approximately 50% of patients achieve durable responses (Sharma and Allison, 2015, Havel et al., 2019). Consequently, there remains a significant unmet need for effective treatment options for malignant melanoma, particularly for patients resistant to standard therapies (Chapman et al., 2011, Flaherty et al., 2012).

Standard cancer treatment like chemotherapy and irradiation have been shown to induce diverse cellular fates in melanoma cells, including cellular senescence (Haferkamp et al., 2013, Mhaidat, 2007, Jost et al., 2021). Cellular senescence was initially observed *in vitro* in primary cells undergoing extensive culture and replicative exhaustion associated with telomere shortening (Hayflick, 1965). It is characterized by a stable proliferation arrest, and therapy-induced senescence has emerged as a common cell fate decision in cancer treatment and a potential target for therapeutic intervention (Chakrabarty et al., 2021, Salama et al., 2014, Fleury et al., 2019). Recent studies have indicated that senescent cells contribute to treatment resistance and the development of secondary cancers in various malignancies, including melanoma (Chakrabarty et al., 2021, Guillon et al., 2019, Thompson et al., 2021).

Evidence from preclinical studies suggests that genetic or pharmacological elimination of senescent cells can attenuate disease progression, improve physical function, and delay all-cause mortality (Baar et al., 2017, Mylonas et al., 2021, Xu et al., 2018, Yousefzadeh et al., 2018). Therefore, targeting senescent cancer cells for elimination using senotherapy (senolytic drugs) holds promising therapeutic potential. Senotherapies may also enhance the efficacy of senescence-inducing cancer treatments by reducing unwanted side effects on overall health (Jochems et al., 2021).

Recently, our team and others have demonstrated the sensitivity of ovarian, breast, sarcoma, prostate, pancreatic ductal adenocarcinoma and glioma cells to senolytics through the selective elimination of senescent cells generated by primary therapies, employing a one-two-punch combinatorial therapeutic approach (Fleury et al., 2019, Lafontaine et al., 2021, Malaquin et al., 2020, Jaber et al., 2023, Balakrishnan et al., 2020). Importantly, these studies have investigated multiple cancer treatment modalities and panels of potential senolytic drugs, revealing the context-dependent nature of senescence induction and senolytic sensitivity. Several studies have provided valuable insights into the mechanisms and implications of senescence in melanoma development, progression, and therapy resistance (Giuliano et al., 2011, Maertens et al., 2013, Thompson et al., 2021). Noteworthy studies have explored the senescence-associated molecular changes, signaling pathways, and interactions with the tumor microenvironment in solid tumors (Oubaha et al., 2016, Milanovic et al., 2018). Furthermore, other studies have investigated the impact of senescence on immunotherapy response and the potential of senolytic drugs to enhance therapeutic outcomes in solid tumors (Ruscetti et al., 2021). Collectively, these studies have underscored the importance of senescence in melanoma as a potential actionable target for therapeutic strategies.

In recent studies focusing on senescence in melanoma, researchers have made significant contributions to our understanding of the implications of senescence in this disease. Liang et al. identified distinct senescence-associated gene expression signatures in melanoma samples, correlating with disease progression and patient outcomes (Liang et al., 2023). Others explored the reciprocal interactions between senescent melanoma cells and the tumor microenvironment, shaping the immune landscape (Lin et al., 2023), and the endothelial senescence signature serving as prognostic markers for survival and immune response prediction (Wu et al., 2023).

Previously, authors demonstrated the potential of targeting the senescence-associated secretory phenotype (SASP) to enhance the response of melanoma cells to immunotherapy (Milanovic et al., 2018). Additionally, others highlighted the role of therapy-induced senescence in promoting drug resistance and found that the presence of senescence markers, such as p16 and SA-β-gal, correlated with poor prognosis and decreased overall survival in melanoma patients (Thompson et al., 2021). These studies collectively underscore the importance of investigating senescence in melanoma and provide valuable insights for developing novel therapeutic strategies.

In this study, our objective was to comprehensively assess the diverse cell fate outcomes induced by clinically relevant therapies in melanoma and investigate the potential of a combination senolytic approach to enhance therapeutic responses in specific contexts. Our findings revealed a wide spectrum of melanoma therapy-induced cell fate decisions, and notably demonstrated that treatments triggering persistent DNA damage in senescent melanoma cells are amenable to a conventional senolytic strategy involving mainly Bcl-2/XL inhibitors.

## 2 MATERIAL AND METHODS

### 2.1 Cell lines and cell culture

Four human melanoma cancer cell lines were included in this study: the Mel-SK23 cell line, which expresses wild-type BRAF, Mel-1102 carrying NRASQ61K mutation, and two cell lines with BRAFV600E mutation (Mel-624.38 and Mel-526). These cell lines were originally obtained from the National Cancer Institute (NCI), NIH, Bethesda, MD USA (Topalian et al., 1989), and were generously provided by Dr. Rejean Lapointe’s Lab at the Centre de Recherche du Centre Hospitalier de l’Université de Montréal (CRCHUM), Canada. Prior to experimentation, all cell lines were routinely screened and confirmed to be negative for mycoplasma contamination. Cells were cultured in RPMI medium with 8% fetal bovine serum, 100 I.U.ml^-1^ penicillin, 100 µg.ml^-1^ streptomycin, and 2mM.ml^-1^ L-glutamine (all obtained from Wisent, QC, Canada).

### 2.2 Viruses and infections

H2B-GFP lentiviruses were produced as described previously (Fleury et al., 2019) and viral titers were adjusted to achieve ∼90% infectivity (Rodier et al., 2011). Infections were followed 48 h later by hygromycin selection (200µg/ml for 3 days) and stable cells were used in experiments.

The generation of H2B-GFP-infected cells and related methodologies can be found in our previous publication (Fleury et al., 2019). Briefly, lentiviruses encoding H2B-GFP were produced using established methods for ∼90% infectivity and 48 hours after infection cells underwent puromycin or hygromycin selection. The lenti-H2B-GFPhygro lentivector was created by amplifying the H2B sequence from the pENTR1A-H2B-HcRed plasmid using specific primers (ES-92 and ES-93). The amplified product was inserted into the entry vector pENTR1A-GFP-N2 (FR1) with Kpn1/BamH1 restriction enzyme digestion before recombining it into the lentiviral destination vector (pLenti PGK Hygro Dest (W530-1)). This final lentivector was used in subsequent experiments.

### 2.3 Drug treatments and irradiation

ABT-263 (Navitoclax) was from APExBIO (Houston, TX, USA). A-1155463 (S7800) was from Selleckchem (Houston, TX, USA). Piperlongumine (1919) was from BioVision (Milpitas, CA, USA). ABT-199 (Venetoclax) and Dabrafenib were from Cayman Chemical (Ann Arbor, MI, USA). Trametinib was from Toronto Research Chemicals (North York, ON, CA). Carboplatin and paclitaxel were from Accord Healthcare (Kirkland, QC, CA). Drugs were first dissolved in 100% dimethyl sulfoxide (DMSO) and then further diluted in complete culture media for each experiment. The drugs were added to the cell culture 24 hours after seeding, and BMi treatment was refreshed every 3 days throughout all experiments. Carboplatin and paclitaxel (10µM/30nM) combination treatment lasted 24 h followed by a wash with PBS (twice) and replenishment of fresh complete culture media. Irradiation was performed using Gammacell® 3000 irradiator Elan at a dose rate of 0.75 Gy/min for a total dose of 10 Gy followed by a fresh media change.

### 2.4 Real-time cell proliferation phase-contrast imaging assay

For live cell proliferation assessment in 96-well, 1000 cells/well were seeded for Mel-SK23, 3000 cells/well were seeded for Mel-1102, and 1500 cells/well were seeded for Mel-624.38 and Mel-526 (all expressing H2B-GFP). Cells were either irradiated (10 Gray) before seeding, or incubated with DMSO (control), or Braf and Mek inhibitors (Dabrafenib+Trametinb at 50/5nM) or carboplatin and paclitaxel (10µM/30nM) at different times. IncuCyte^™^ Live-Cell Imaging System (IncuCyte HD) was used to image cell number by phase contrast and fluorescence. Frames were captured every 8 hours (10X objective). Proliferation data were analyzed by using IncuCyte^™^ S3 software based on green element count (H2B-GFP cell nuclei) or cell confluency and growth curves were plotted by GraphPad Prism 9.0 software (GraphPad Inc., San Diego, CA). Each experiment was performed in triplicate and repeated three times.

### 2.5 Real-time cell death imaging and specific death assays

Propidium iodide (PI) fluorescently label dead cells based on the loss of cell membrane integrity (PI is normally a cell impermeant DNA binding dye (Belloc et al., 1994). PI (Sigma Aldrich, Saint-Louis, USA) was added to the culture media at a concentration of 0,5µg/ml, and cells imaged as for the real-time proliferation assay. Frames were captured at 8h intervals from two separate regions per well using 10X objective. The IncuCyte^™^ S3 software was used to quantify the percent of dead cells by scoring PI positive fluorescently labeled red nuclei against total green H2B-GFP cell nuclei. Graph were plotted using GraphPad Prism 9 (GraphPad Inc., San Diego, CA). To assess specific cell death caused by senolytic combinations we further normalized the death data using 2 strategies, first by removing the baseline level of cell death in each condition (from the matched control), second by creating a 100% death measurement using localized cell lysis with a short Triton treatment at concentration of 0,01% in the culture media (TritonTM X-100 solution, Sigma Aldrich, Saint-Louis, USA) (Cummings and Schnellmann, 2004). To further ascertain that we measured the additional cell death caused by the senolytic treatment, prior to senolytics drugs and PI addition, melanoma cells (treated or untreated) were washed with PBS to remove floating cells. Each experimental condition was performed in triplicate and repeated at least three times. For quantification of specific death we adapted the chromium-51 release assay formula (Wallace et al., 2004):

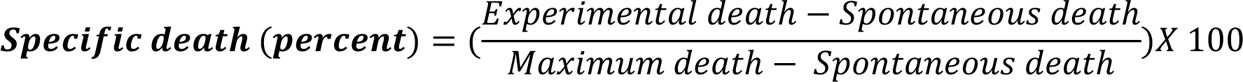

Experimental death = (PI+/H2B-GFP+) / (H2B-GFP+) In each experimental condition

Spontaneous death = (PI+/H2B-GFP+) / (H2B-GFP+) In control (DMSO) condition

Maximum death = (PI+/H2B-GFP+) / (H2B-GFP+) In Triton treated condition

Each experiment was performed in triplicate and repeated at least 2 times.

### 2.6 Clonogenic assays

Cells were seeded in a 6-well plate at a density of 500 cells (Mel-SK23), 2000 cells (Mel-1102) or 1000 cells (Mel-624.38 and Mel-526) per well. The media was removed and replaced with complete media (RPMI 8%FBS) containing Braf and Mek inhibitors alone or in combination at indicated concentrations. Cells were treated for 15 days and analyzed or released for an additional 15 days in a drug-free medium and analyzed. Cells were fixed and colored with a mix of 50% v/v methanol and 0.5% m/v of crystal violet (Sigma-Aldrich Inc., St. Louis, MO). Colonies were counted under a stereomicroscope at a ×2 magnification and reported as a percentage of control. Each experiment was performed in duplicate and repeated 2 times.

### 2.7 Immunofluorescence and pulsed DNA synthesis detection

Cells were seeded in 8-wells chamber slides (Life Sciences, Corning, NY, USA) and allowed to adhere for 24 h before exposition to treatments. Cells were fixed for 5 min in formalin at room temperature (RT) and permeabilized in 0.25% Triton in phosphate-buffered saline (PBS) for 10 min. Slides were blocked for 1 h in PBS containing 1% bovine serum albumin (BSA) and 4% donkey serum. Primary antibodies (γH2AX and 53BP1) diluted (1/2500) in blocking buffer were added in each well and slides were incubated overnight at 4 °C. Cells were washed (PBS) and incubated with secondary antibodies (dilution 1/5000) for 1 h at RT, then washed again.

To detect DNA synthesis EdU (5-ethynyl-2′-deoxyuridine; 10 μM, Invitrogen) was added to the medium and incubated for 24 h from days 8 to 9 post-treatment. Cells were washed three times with TBS and fixed with 10% formalin for 5 min. EdU fluorescence staining was assessed using the Click-iT^®^ EdU Alexa Fluor^®^ 488 Imaging Kit (Invitrogen).

For immunofluorescence and EDU coverslips were mounted onto slides using Prolong^®^ Gold antifade reagent with DAPI (Life Technologies Inc.). Images were obtained using a Zeiss microscope (Zeiss AxioObserver Z1, Carl Zeiss, Jena, Germany). An automated analysis software from Zeiss (AxioVision^™^, Carl Zeiss) was used to count DNA damage foci to calculate the average number of foci per nucleus. The fold change was calculated as the ratio between percentages of γH2AX or 53BP1 nuclear foci in treated versus control (nontreated) cells. DNA damage positive cells percentage was calculated relative to the DAPI staining (total nuclei count). γH2AX and 53BP1 foci were quantified in more than 50 nuclei from 3 different fields of each chamber. For DNA synthesis evaluation EdU positive cells were similarly counted and reported to total nuclei (DAPI).

### 2.8 Analysis of cell cycle by flow cytometry

Cells were seeded in 6-well plates and treated 24 h after seeding. At each indicated time, cells were trypsinised, washed with PBS and fixed in cold ethanol (70%) for 24 h. Cells were then washed with PBS and stained for 30 min at room temperature with a 25 μg/ml PI solution containing 100 μg/ml RNAse A. The PI fluorescence signal was detected using the Fortessa flow cytometer (BD Biosciences, Mississauga, ON) and analyzed with FlowJo™ v10.8 Software (BD Life Sciences).

### 2.9 SA β-galactosidase detection

We adapted the SA β-galactosidase protocol used by Fleury et al (Debacq-Chainiaux et al., 2009, Fleury et al., 2019). Briefly, cells were seeded in 6-well plates and treated 24 h after seeding. At the endpoint (day 9 post-treatment), cells were washed once with PBS, fixed with 10% formalin for 5 min, washed again with PBS, then incubated at 37 °C for 12–16 h (depending on the cell line) in a staining solution composed of 1 mg ml^−1^ 5-bromo-4-chloro-3-inolyl-β-galactosidase in DMSO (20 mg ml^−1^ stock), 5 mM potassium ferricyanide, 150 mM NaCl, 40 mM citric acid/sodium phosphate, and 2 mM MgCl_2_, at pH 6.0. Finally, cells were washed twice with PBS and at least 4 representative pictures per well were taken for quantification using EVOS™ FL Digital Inverted Fluorescence Microscope from Thermo Fisher Scientific (Carlsbad, CA, USA). Each experiment was performed in duplicate (6 well plate) and repeated 3 times.

### 2.10 Protein extraction and western blot analysis

Cells were seeded in petri dishes (100 mm) and allowed to adhere for 24h before each treatment. At the indicated times, cells were lysed in mammalian protein extraction reagent (MPER, Thermo Fisher Scientific, Waltham, MA) containing a protease and phosphatase inhibitor cocktail (Sigma-Aldrich Inc., St. Louis, MO). After protein quantification (Pierce BCA Protein Assay Kit, Thermo Fisher Scientific), 15 micrograms of total protein were separated using stain-free 4–15% gradient Tris-glycine SDS-polyacrylamide gels (Mini PROTEAN^®^ TGX Stain-Free^™^ Gels, Bio-Rad Laboratories, CA) and transferred onto PVDF membranes (Amersham Hybond, GE Healthcare Life Sciences, Mississauga, ON, Canada). Immunodetections were performed using enhanced chemiluminescence (Thermo Fisher Scientific) to detect peroxidase-conjugated secondary antibodies bound to primary antibodies. A ChemiDoc MP Imaging System (Bio-Rad Laboratories) was used to detect chemiluminescence. The stain-free technology (Bio-Rad Laboratories) was used to quantify protein loading in the gel/membrane. Immunoreactive band intensities were quantified using ImageJ software (https://imagej.nih.gov/ij/).

### 2.11 IL8 secretion measurement in conditioned medium

Conditioned media (Maertens et al.) were prepared by incubating cells with RPMI complete medium (FBS 8%) for 48 h and stored at −80 °C until probed. Levels of IL-8 were assessed using ELISA (R&D Systems (IL-8 #DY208). The data were normalized to cell number and reported as fold change of secreted protein compared to control.

### 2.12 Antibodies

The following antibodies were used: phospho-histone γ-H2AX (clone JBW301, EMD Millipore, Temecula, CA) (dilution for immunofluorescence 1/2500); 53BP1 (clone 305, Novus Biologicals, Littleton) (dilution for immunofluorescence 1/2500). p53 (clone DO-1, Santa Cruz 1/5000) ERK-1/2 and phospho-ERK-1/2, 1/2500, Cell Signaling, #4370S; phospho-Rb (Ser 807/811), Cell Signaling, 1/2500; phospho-p90-RSK, Cell Signaling, 1/2000; p21, BD, 1/2000; GAPDH, Cell Signaling, 1/2500; Tubulin-a, 1/5000, Cell Signaling, 3873P).

## 3 Results

### 3.1 Varied melanoma treatments trigger distinct cell fate phenotypes

To investigate the effects of various cancer therapies on melanoma cell fate decisions, we selected a panel of four human melanoma cell lines with distinct genomic backgrounds (Fig 1. A left) that reflect clinical features of malignant melanoma: Mel-SK23 (wild-type BRAF), Mel-1102 (NRASQ61K mutation), Mel-624.38 and Mel-526 (BRAFV600E and TP53 mutations). Lentiviruses encoding H2B-GFP were used to transduce all cells, enabling stable labeling of cell nuclei with GFP fluorescence. To mimic targeted melanoma therapy, we initially exposed these cells to increasing concentrations of Braf and Mek inhibitors alone or in combination (Dabrafenib+Trametinib, referred to as BMi). Using live cell imaging (Incucyte), we monitored cell proliferation following the treatments (Fig S1. C-E). As anticipated, Dabrafenib alone selectively inhibited the proliferation of Braf V600E mutant melanoma cells (Mel-624.38 and Mel-526) in a dose-dependent manner, while wild-type Braf Mel-SK23 or NRASQ61K Mel-1102 cells maintained their proliferative potential (Fig S1.C).

**Figure 1:**
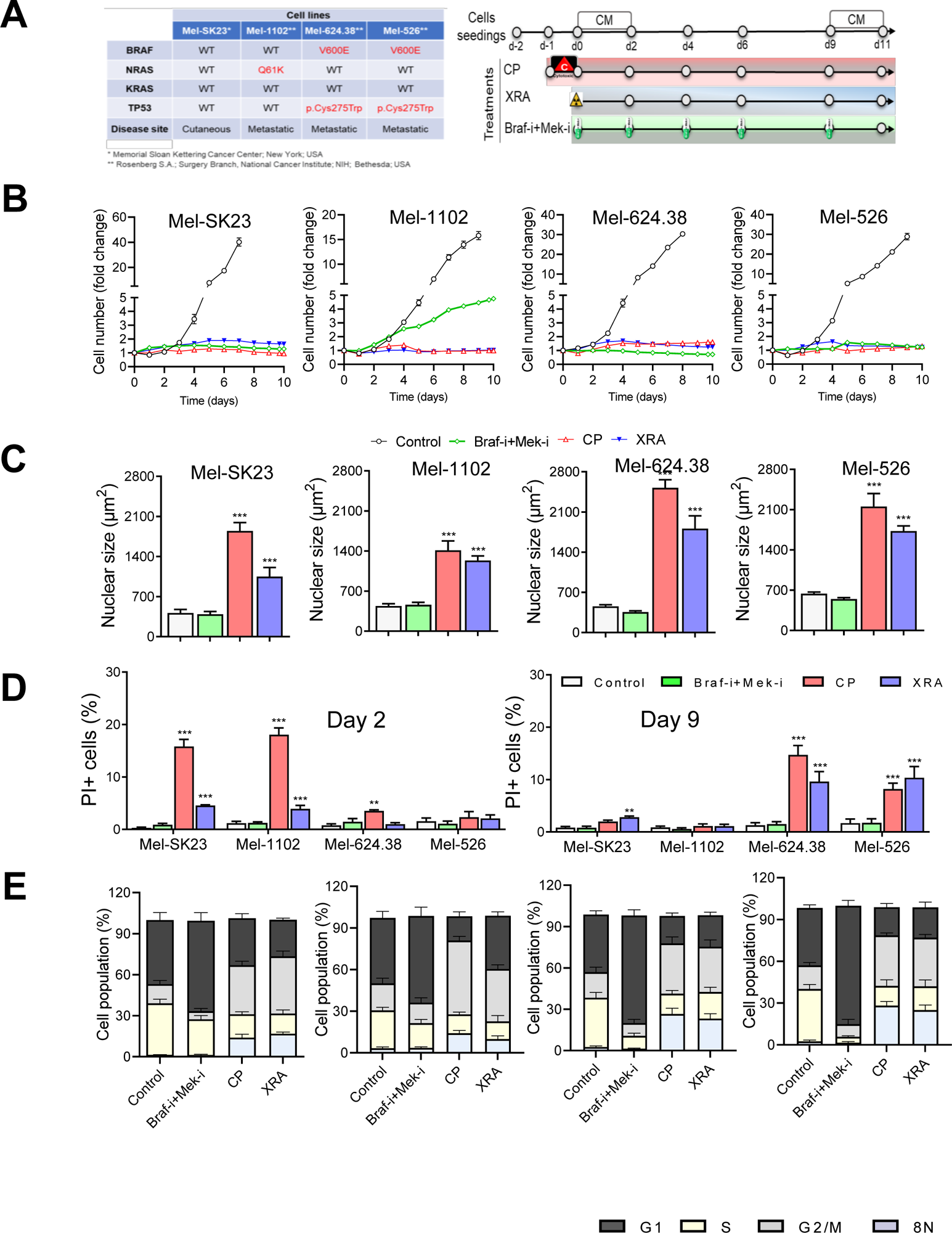
Genotoxic treatment trigger cell death with distinct phenotype compared to BMi in melanoma cells. **A)** Table showing the genetic background of the panel of selected melanoma cell lines (left) and treatment timeline (right). **B)** Representative proliferation curves of melanoma cells exposed to different treatments (Dabrafenib – Braf-i (50nM) + Trametinib – Mek-i (5 nM, Braf-i+Mek-i), Carboplatin plus Paclitaxel - CP (10 µM + 30 nM) or Radiation - XRA (10 Gray). **C)** Average nuclear size of melanoma cells expressing H2B-GFP (untreated or day 6 post-treatment). **D**) Cell death quantification at by PI incorporation at day 2 or day 9 post-treatment through live imaging. **E**) Cell cycle distribution of 9 days treated melanoma cells compared to their controls. Mean ± SD of 2-3 independent experiments is shown. Student’s t-test, * *p* < 0.05, ** *p* < 0.01, *** *p* < 0.001.

In contrast, Trametinib alone inhibited the proliferation of all four cell lines, although it was more effective in Mel-624.38 and Mel-526 cells (Fig S1.D). The combination of both inhibitors exhibited striking efficacy, completely inhibiting proliferation in Braf mutant cells, while NRASQ61K Mel-1102 cells showed reduced proliferation at the lowest doses of the combination (Fig 1.B and S1.E). Consistently, phosphorylation of ERK and its downstream effector p90-RSK were undetectable after 48 hours of inhibition in Mel-SK23, Mel-624.38, and Mel-526 cells, whereas Mel-1102 cells displayed low levels of ERK and p90-RSK phosphorylation (Fig S1.B). Thus, BMi effectively block MAPK signaling, except in Mel-1102 cells, which may explain their retained proliferation. That is consistent with the limited efficacy of these inhibitors in patients harboring NRASQ61K mutation (Dummer et al., 2017). Mel-1102 cells might have a more prominent dependence on the MAPK pathway (for survival and proliferation), which is effectively targeted by Trametinib (Sun et al., 2014). However, when both drugs are combined, there could be a compensatory mechanism where the inhibition of BRAF by Dabrafenib leads to an increased activation of MEK (through CRAF), which might lead to reduced growth inhibition. This is consistent with the observed phosphorylation of p90-RSK, a downstream effector of ERK, 48 hours following BMi treatment in Mel-1102 cells (Fig S1.B). Overall, these findings validate the genetic profile of MAP-kinase signaling in our melanoma cell line panel.

Additionally, we treated the panel of melanoma cells with conventional chemotherapies and radiotherapies, which are occasionally used for melanoma treatment in clinical settings (Kuryk et al., 2020). Cells were exposed to a combination therapy of carboplatin plus paclitaxel (10 µM of Carboplatin + 30 nM of Paclitaxel, referred to as CP), or to a single dose of ionizing radiation (10 Gy of X-ray, referred to as XRA), previously demonstrated to impede cancer cell proliferation and to induce stable senescence in human normal and cancer cells (Coppe et al., 2008, Malaquin et al., 2020, Rodier et al., 2009). Notably, we observed nearly complete proliferation arrest in all melanoma cells following either CP or XRA treatment, irrespective of their Braf or other mutation statuses (Fig 1.B).

Cancer cells can undergo diverse fates in response to anti-cancer therapies, including cell death, senescence, mitotic catastrophe, autophagy or altered proliferation (Fu et al., 2021). Thus, we evaluated which cell fate was induced by BMi, CP, or XRA in melanoma cells. Live cell imaging revealed that CP and XRA treatments led to nuclei enlargement within 2 days, a feature absent even in long-term BMi-treated cells (Fig 1.C, S2.A). Nuclei enlargement often signifies increased ploidy and genome instability (Storchova and Pellman, 2004). Therefore, we assessed cell cycle distribution using flow cytometry 9 days after each treatment and observed G1 phase accumulation in BMi-treated cells, while CP and XRA induced G2/M accumulation, accompanied by a significant increase in 8N polyploid cells (Fig 1.E). Raw flow cytometry profiles also suggested a sub-G1 subpopulation in CP and XRA, but not in BMi (Fig S1.F), indicating genomic instability and cell death in response to DNA-damaging agents.

Subsequently, we examined whether the treated melanoma cells underwent cell death. To assess this, we employed a real-time imaging assay based on dynamic propidium iodide (PI) incorporation. Our observations revealed that CP and XRA treatments induced cell death in melanoma cells, whereas no cell death was observed upon treatment with the combination of BMi (Fig 1.D). Interestingly, the cell death triggered by CP and XRA primarily occurred earlier in wild-type p53 cells (Mel-Sk23 and Mel-1102), while it occurred later in mutated p53 cells (Mel-624.38 and Mel-526).

### 3.2 Figure 2: Surviving melanoma cells exhibit a senescence phenotype following DNA damaging treatment

We observed that while some melanoma cells treated with CP and XRA underwent cell death, the surviving cells displayed a senescent phenotype. Given the central role of cellular senescence in response to cancer therapies (Fleury et al., 2019, Malaquin et al., 2020), we further assessed senescence hallmarks in surviving melanoma cells.

**Figure 2:**
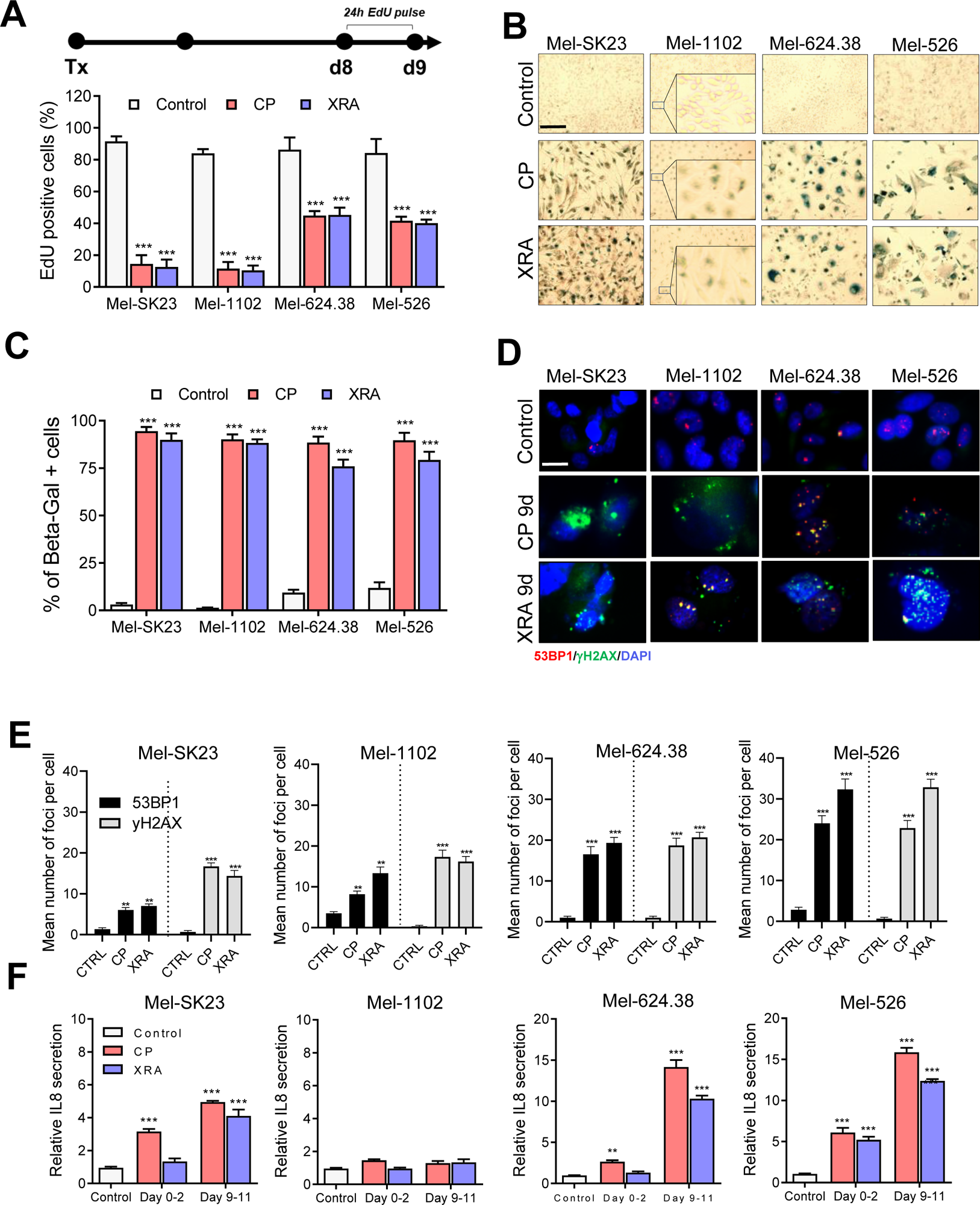
Surviving melanoma cells exhibit a senescence phenotype following genotoxic treatment. **A)** Timeline of EdU pulse (top) and quantification (bottom) of 24 hours EdU positive cells from day 8 to 9 post-treatment. **(B, C)**: Representative images **B)** and quantification **C)** of SA-β-gal staining in melanoma cells fixed 9 days after C/P or XRA treatments. (**D, E)**: Representative images **D)** and quantification **E)** of γH2AX and 53BP1 foci in indicated cell lines by immunofluorescence. **F)** Quantification of IL8 secretion measured by ELISA from day 0-2 or day 9-11 post-treatments in the indicated cells. Mean ± SD of 3 experiments is shown. Student’s t-test, * p < 0.05, ** p < 0.01, *** p < 0.001.

To evaluate the proliferative capacity of surviving cells we conducted a 24-hour Edu pulse assay, which measure DNA synthesis (Fig 2.A top). Nine days post-CP and XRA treatments, we observed a significant reduction in Edu-positive cells in the Mel-SK23 and Mel-1102 cell lines (about 20% residual positivity), whereas approximately 40% of cells in the Mel-624.38 and Mel-526 lines remained Edu-positive (Fig 2.A, Fig S2.B). This data, along with cell cycle distribution analysis, reveal that CP and XRA treatments resulted in a G2/M phase arrest in melanoma cells following S phase, with more pronounced aneuploidy (8N) and genome instability in p53-mutated melanoma cells compared to wild-type p53 cells.

Additionally, SA-β-Gal staining revealed a significant increase in beta-galactosidase activity in melanoma cells nine days after treatment with CP and XRA (Fig 2.B, C), while immunofluorescence assays performed nine days post-treatment, demonstrated the presence of persistent DNA double-strand break damage foci decorated by 53BP1 and phosphorylated histone H2AX (Fig 2.D, E). Additionally, these senescent cells exhibited elevated levels of p53 and p21 proteins (Fig S2.C-F).

Accumulation of DNA damage response (DDR) foci and genome instability have been shown to contribute to the induction of cellular senescence (Ghadaouia et al., 2021). Activation of NF-kappaB, which is essential for upregulation of SASP cytokines (Elliott et al., 2001, Wang et al., 2017), such as interleukin 6 (IL6) and interleukin 8 (IL8), is another consequence of DDR foci accumulation and genome instability. Although we did not detect IL6 secretion by ELISA in these melasnoma cells, we assessed IL8 and observed a time-dependent significant increase in IL8 secretion in the treated cells, except for the Mel-1102 cell line (NRASQ61K), which showed no significant difference in IL8 secretion over time. Compared to untreated cells, the CP and XRA treated cells exhibited 4 to more than 10 times higher levels of IL8 secretion (Figure 2.F). Collectively, these findings demonstrate that melanoma cells exhibit a senescence phenotype in response to DNA damage-inducing treatments with CP and XRA.

### 3.3 Long-term BMi triggers a partially reversible senescence-like phenotype in Braf V600E melanoma cells

Next, to assess senescence hallmarks following combined Braf and Mek inhibition, we continuously exposed melanoma cells to BMi for 9 days. In a 24-hour EdU pulse experiment (Fig 3.A top), consistent with the dose-response growth curve (Fig S1.E), we observed that wild-type Braf melanoma cells (Mel-SK23 and Mel-1102) had a relatively high frequency of EdU-positive cells (19% and 50% respectively), while Braf V600E mutated melanoma cells (Mel-624.38 and Mel-526) showed almost no EdU incorporation (Fig3.A bottom). This suggests that the inhibitors induced a more robust proliferation arrest in Braf mutated cell lines. Importantly, consistent with previous findings (Krayem et al., 2018), we observed an increase in SA-Beta-galactosidase activity in Braf V600E mutated cells 9 days after treatment with BMi, but not in wild-type Braf cells (Fig3. B, C).

**Figure 3:**
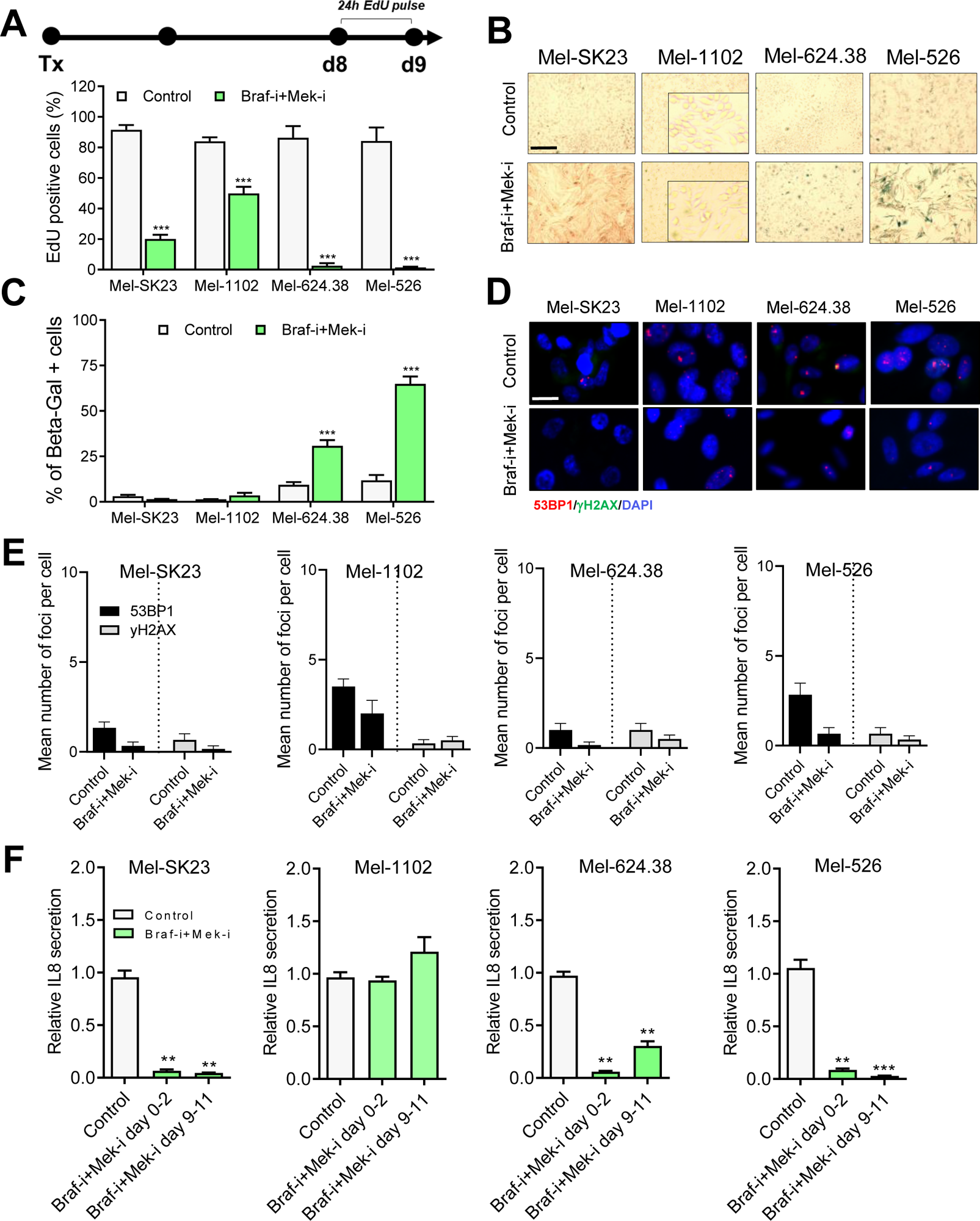
BMi trigger a senescence-like phenotype in Braf V600E melanoma cells. **A)** Timeline of EdU pulse (top) and Quantification (bottom) of 24 hours EdU positive cells from day 8 to 9 post BMi treatment. **(B, C)** Representative images **B)** and quantification **C)** of SA-β-gal staining in melanoma cells fixed 9 days after BMi treatment. **(D, E)** Representative images **D)** and quantification **E)** of γH2AX and 53BP1 foci in indicated cell lines by immunofluorescence. **F)** Quantification of IL8 secretion measured by ELISA from day 0-2 or from day 9-11 following BMi treatment in the indicated cells. Mean ± SD of 3 experiments is shown. Student’s t-test, * p < 0.05, ** p < 0.01, *** p < 0.001.

However, unlike CP and XRA treatments, BMi treated cells did not show increased DDR foci (Fig3.D, E). Correspondingly, BMi treated cells did not increase their secretion of IL8, which correlates with the absence of DDR foci (Fig3.F). The distinct phenotype observed in BMi treated cells supports the idea that the response of cancer cells to therapy depends not only on the cell type but also on the type of treatment. Taken together, these data suggest that long-term Braf and Mek inhibition triggers a partial senescence-like phenotype in melanoma cells harboring the Braf V600E mutation.

Although senescence was initially defined as an irreversible proliferation arrest, recent studies have shown that some TIS cancer cells can escape the senescence state and re-enter the cell cycle (Flaherty et al.). For example we previously reported that Olaparib-induced senescence in high-grade serous ovarian cancer cells or androgen depletion-induced senescence in prostate cancer are reversible (Fleury et al., 2019)(MALAQUIN 2020). Therefore, we investigated whether the senescence-like state induced by BMi is stable or primarily due to sustained inhibition. To address this, we performed a real-time imaging proliferation assay during and after BMi treatment at different time points (Fig4.A). We observed that wild-type Braf cells quickly recovered from the growth inhibition and reached confluency after 9 or 15 days of treatment. However, Braf V600E mutated cells had a pronounced delay in recovering from growth inhibition compared to wild-type cells (Fig 4.B).

**Figure 4:**
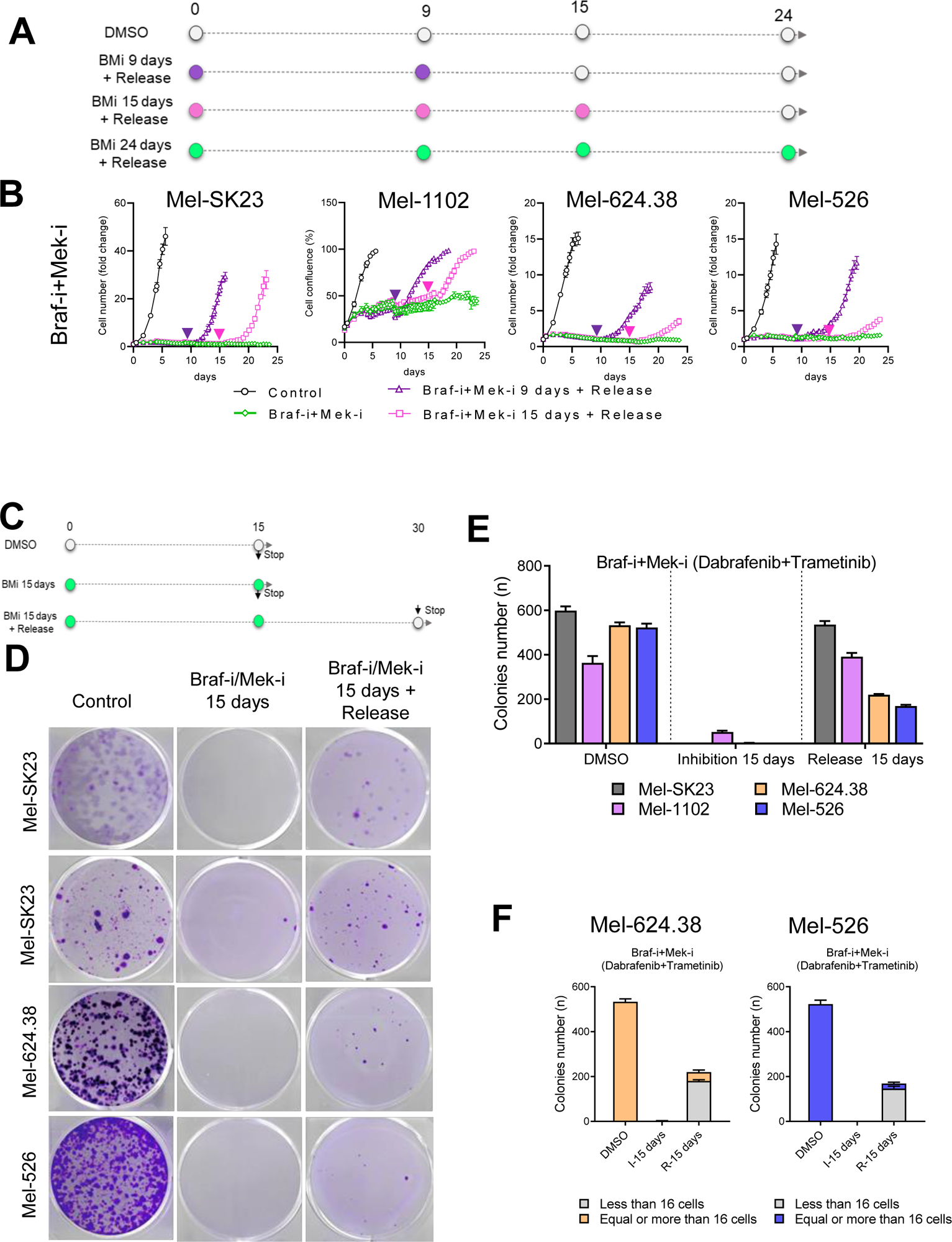
BMi-induced senescence-like phenotype in melanoma cells is partially reversible. **A)** Experimental schema and **B)** proliferation curves of BMi treatment with or without release at the indicated time point in four different melanoma cell lines. **C)** Colonies formation assay timeline and **E)** representative images of 15 days following BMi treatment, or 15 days of BMi treatment and 15 days of release. **E)** Histogram of colonies number from images shown in D. **F)** Histogram of colonies number from images shown in D categorized in two sizes groups (less than 16 cells and more than 16 cells). Data are the mean +/- SD of triplicate and are representatives of three independent experiments.

Furthermore, to determine if the recovery occurred in most cells or only in a proportion of cells, we performed a colony formation assay with BMi for 15 days of treatment followed by an additional 15 days of release without drugs (Fig 4. C). We found that most of the wild-type Braf melanoma cells regained their proliferative capacity after drug release (Fig 4. D, E). In contrast among Braf mutated cells, only around 30% within the treated populations regained their proliferative capacity and formed small colonies mostly composed of 16 cells or less (Fig 4. F). The remaining cells were unable to proliferate even long time after drug release. This suggests the occurrence of possible epigenetic modifications (Crouch et al., 2022) following long-term dual Braf and Mek inhibition, which likely convert Braf V600E melanoma cells into a stable growth arrest state resembling senescence.

Taken together, these data demonstrate that long-term dual Braf and Mek inhibition induces a partially reversible senescence-like state in Braf V600E melanoma cells. Considering that CP and XRA induce a stable senescence phenotype and that Braf V600E cells were also partially converted into a senescence-like state following BMi treatment, we investigated their sensitivity to known senolytics.

### 3.4 DNA damage-induced senescence promotes melanoma cell sensitivity to Bcl-XL/Bcl2 inhibitors

Senescent cells exhibit various characteristics, including senescence-associated apoptosis resistance (SAAR). Identification and understanding of key factors associated with SAAR have led to the development of pharmacological approaches for selective clearance of senescent cells (Baker et al., 2011, Chang et al., 2016). Senolytics are defined as small molecules that can selectively or preferentially eliminate senescent cells (Chang et al., 2016, Hickson et al., 2019, Zhu et al., 2016) without or with minimal cytotoxicity on proliferating or non-senescent cells at a given dose. Recent studies have shown that sensitivity to senolytics, particularly Bcl inhibitors, can vary depending on the type of cancer and the senescence inducer (Fleury et al., 2019, Lafontaine et al., 2021, Malaquin et al., 2020). Given the distinct phenotypes observed in melanoma cells treated with CP, XRA, or BMi, we investigated their sensitivity to a panel of known senolytics.

To assess cell sensitivity to senolytic drugs, we used a novel real-time imaging death assay that capture cell death induced within 24 hours of senolytic treatment. We selected four well-known senolytic drugs: ABT-263, Bcl-xl and Bcl-2 inhibitor (Wendt, 2008), A-115 (Bcl-xl inhibitor) (Tao et al., 2014), ABT-199 (Bcl-2 inhibitor) (Chang et al., 2016), and Piperlongumine (PPL, a natural antioxidant product) (Wang et al., 2016). As expected, we found that CP or XRA induced senescent cells showed significant sensitivity to the selected senolytics, with some variations (Fig 5. B-E, right). ABT-263 and A-115 were generally more effective than ABT-199 in inducing cell death of TIS melanoma cells. In this context, PPL displayed the lowest sensitivity, significantly different from the other drugs. These results showed that both Bcl-xl and Bcl-2 contribute to the survival of CP or XRA-induced senescent melanoma cells.

**Figure 5:**
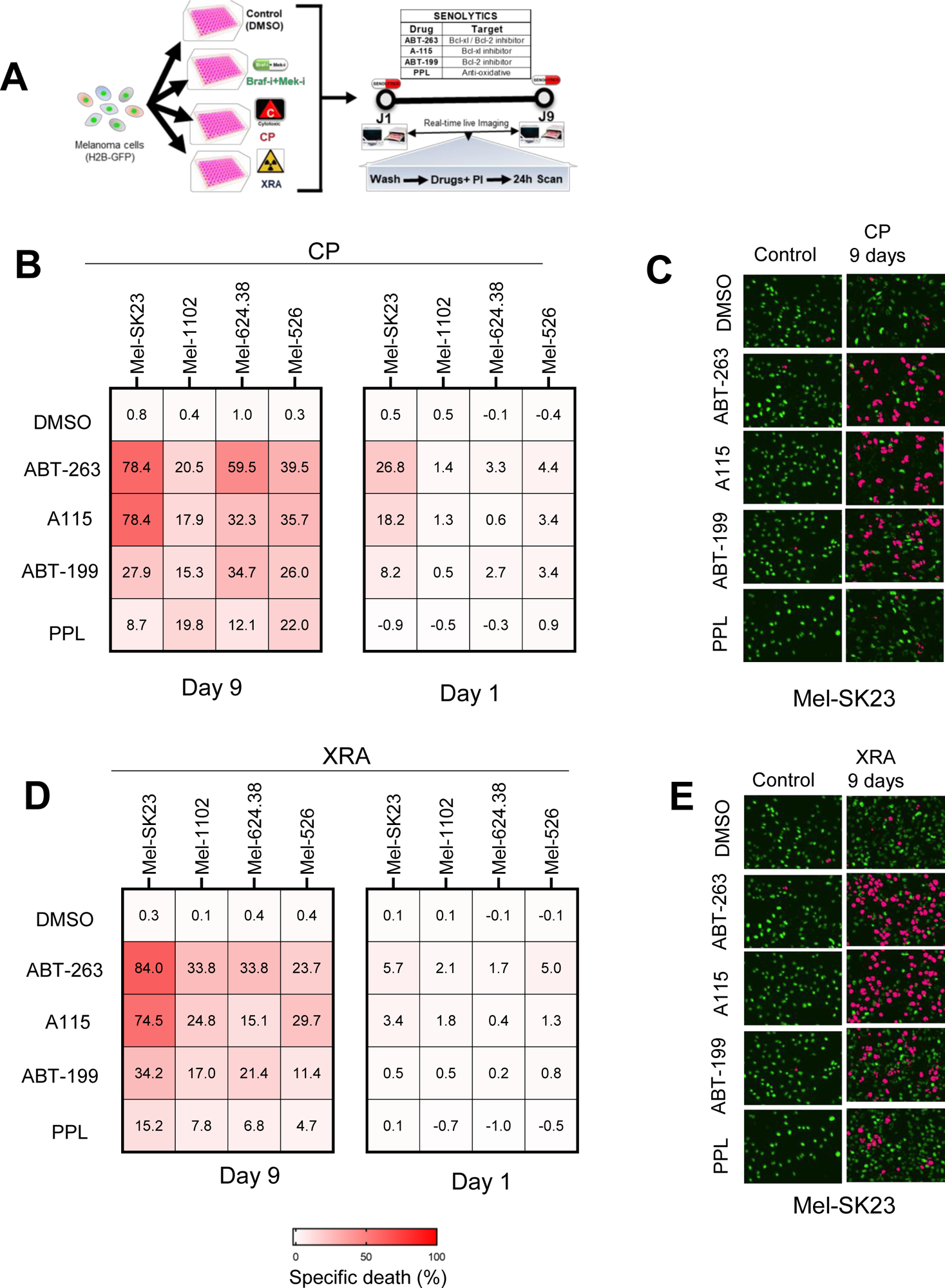
DNA damage-induced senescence but not BMi-induced senescence sensitize melanoma cells to Bcl-xl / Bcl-2 inhibitors. **A)** Experimental design of senolytic assay. **(B, C)** Heat map showing percent of specific death **B**) and representative images **C**) 24 hours following treatment with senolytics (ABT263 - 0.32µM; A115 - 0,32 µM; ABT199 1.25 µM; or PPL - 0.32µM)) in the four melanoma cells, initially pre-treated with C/P for 1 or 9 days. **(D, E)** Heat map showing percent of specific death **D**) and representative images **E)** 24 hours following treatment with senolytics (ABT263 - 0.32µM; A115 - 0,32 µM; ABT199 1.25 µM; or PPL - 0.32µM)) in the four melanoma cells, initially exposed to XRA for 1- or 9-days. Percentage of specific death were calculated using chromium-51 assay release. Data were analyzed using two-tail Student t test. * p < 0.05, **p < 0.01, and ***p < 0.001.

To validate that this sensitivity is due to a senolytic effect, we repeated the same assay one day after CP or XRA treatment (Fig 5.B-E, left). We observed that 3 out of 4 cell lines were not sensitive to these drugs, one day following CP or XRA treatment, confirming that the sensitivity of CP or XRA-treated cells is mainly driven by the senescence state rather than an additive therapeutic effect. However, we noticed that Mel-SK23 cells were sensitive to Bcl-xl/Bcl-2 inhibitors just one day after exposure to CP or XRA. Since both CP and XRA are genotoxic agents, this early sensitivity suggests that not only the senescence state but also the unresolved DNA damage response (DDR) foci (compromising state to senescence) could be sufficient to sensitize Mel-SK23 cells to Bcl-xl and Bcl-2 inhibitors. Taken together, these data demonstrate that Bcl-xl and Bcl-2 family inhibitors can be used as senolytics to provoke cell death in DNA damage-induced senescent melanoma cells.

### 3.5 Direct synergy between combo BMi and Bcl xl / Bcl-2 inhibitors promote melanoma cells death in TIS independent manner

We investigated whether BMi-induced senescent-like cells showed similar senolytic sensitivity. However, due to the synthetic lethality between Mek and Bcl-2/XL inhibitors (Corcoran et al., 2013, Cragg et al., 2008, Iavarone et al., 2019) and the partially reversible nature of the induced phenotype when BMi are discontinued (Fig 4.B), we performed the senolytic assay using a sequential treatment approach, with or without BMi. This approach allowed us to differentiate the effects attributable to the induced senescence-like state from the effects resulting from a direct synergy between Bcl-2/XL inhibitors and BMi.

In contrast to the observed sensitivity of CP and XRA-induced senescent cells, we found that BMi-induced senescent-like cells were not sensitive to Bcl-XL/Bcl-2 inhibitors or to PPL in the sequential combo treatment approach (no significant cell death compared to control cells - Fig 6.B left). However, when the senolytics were added simultaneously with BMi, we observed a significant increase in cell death, particularly with Bcl-XL/Bcl-2 inhibitors (Fig 6. B right). This suggests that, in this context, a direct synergy between Bcl-2/Bcl-XL inhibitors and BMi, rather than the BMi-induced senescence-like state, promotes melanoma cell death (Airiau et al., 2016).

**Figure 6:**
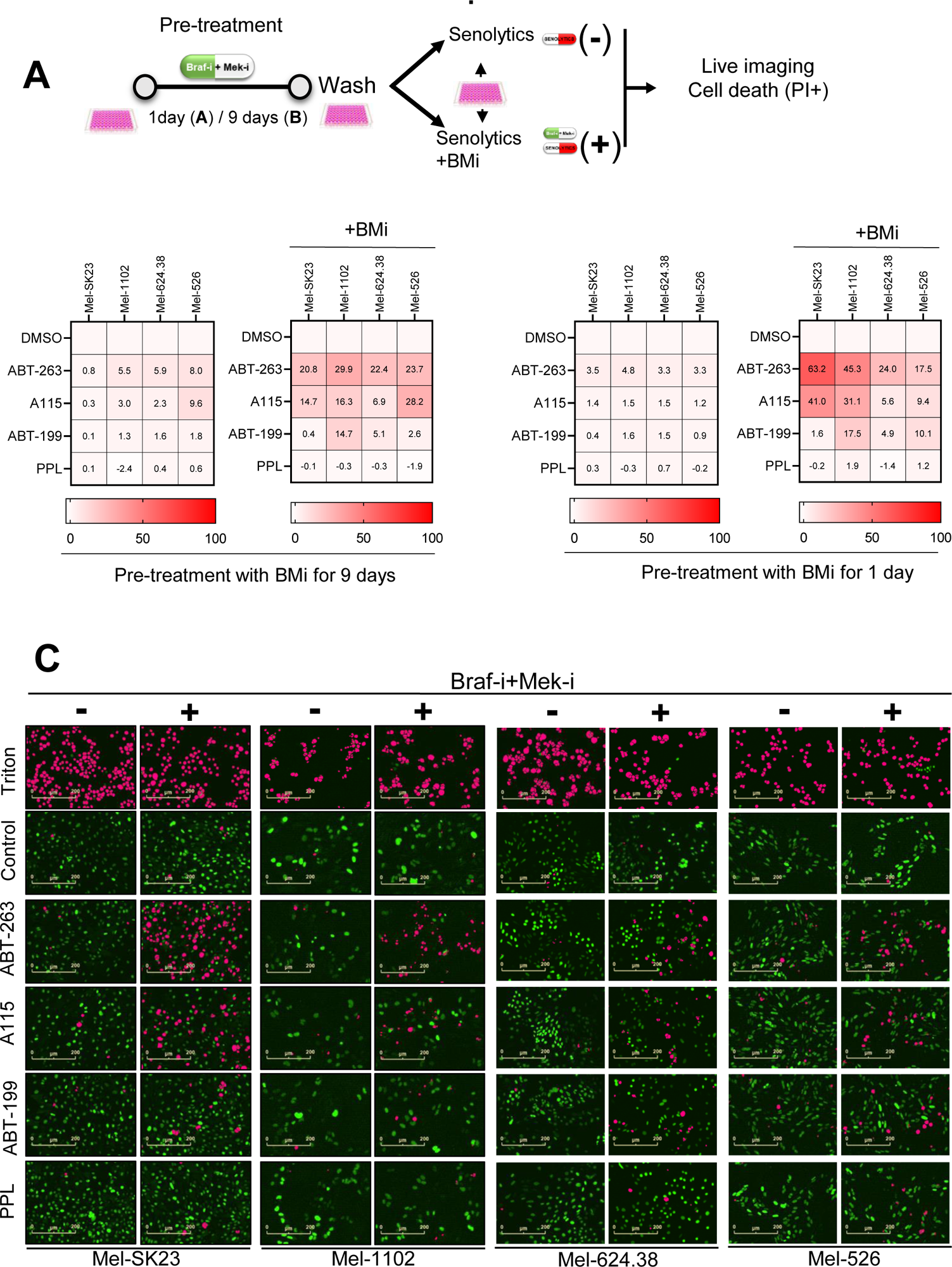
Direct synergy between BMi and Bcl xl / Bcl-2 inhibitors promote melanoma cells death in a cell fate independent manner. **A)** Experimental design of drug synergy assessment based on the specific death. Prior to the synergy assessment, cells were pretreated first with BMi for 1 or 9 days. **B)** Heat map showing percent of specific death 24 hours following treatment with senolytics alone (-) or simultaneously with BMi (+) in different melanoma cells lines, pretreated with BMi for 1 day (top) or 9 days (bottom). **C)** Representative images of data shown in B (H2BGFP in green; and mask of PI/H2B-GFP double positive in pink).

To further confirm that the observed cell death is due to a synergistic effect, we conducted another real-time imaging cell death assay 1 day following BMi treatment (Fig 6. C). We added each senolytic compound with or without BMi. Similarly, we observed that the simultaneous combination of Bcl-2/Bcl-XL inhibitors and BMi provoked melanoma cell death (Fig 6. C right), which was not the case when we used Bcl-2/Bcl-XL inhibitors alone (Fig 6. C left).

Overall, these results demonstrate that, by using the one-two punch approach, BMi-induced senescence-like phenotype does not sensitize melanoma cells to Bcl-2/Bcl-XL inhibitors and PPL. Bcl-2/Bcl-XL inhibitors need to directly cooperate with BMi to promote melanoma cell death.

## 4 Discussion

In this study, we investigated the effects of different therapies on melanoma cell fate decisions, with a specific focus on therapy induced senescence and the potential for combination treatments with senolytic drugs. Cellular senescence is considered as a double edged sword as it is a potentially beneficial cell fate decision in response to most cancer treatments given its effect on the proliferation of damaged cells, but the presence of senescent cells has also been shown to promote the proliferation of nearby non-senescent cells (Sun et al., 2018). Accordingly, senescence has been associated with resistance and the development of secondary cancers (Chakrabarty et al., 2021, Guillon et al., 2019, Thompson et al., 2021). Our results provide valuable insights into the response of melanoma cells to clinically relevant therapies including CP, XRA and BMi targeted therapies and reveal the diversity of senescent phenotypes produced, which have implications for the design of more effective treatment strategies against melanoma.

As expected, BMi effectively blocked MAP-kinase signaling and selectively inhibited the growth of Braf mutant melanoma cells. This is consistent with the clinical response observed in patients harboring Braf mutations, highlighting the importance of personalized treatment approaches based on the genetic profile of the tumor. Notably, we observed that NRAS-mutated melanoma cells showed reduced proliferation upon treatment with the combination of BMi, suggesting some level of sensitivity to these targeted therapies. However, the limited efficacy in NRAS-mutated cells underscores the need for alternative treatment options for this subset of melanoma patients.

Furthermore, we observed that CP and XRA treatments resulted in a mixed response of cell death and cellular senescence, regardless of BRAF mutation profiles. This observation is consistent with previous studies showing that cellular senescence is a common responses of melanoma cells to CDDP-based treatment (Sun et al., 2018). The authors showed that CDDP-induced senescence through DDR and p53/p21 axis. Since p21 is direct p53 target, its expression in p53 mutant cells could be explained by the involvement of other mediators such as EZH2 (Flaherty et al.). EZH2 is a transcriptional repressor whose expression is correlated with the malignant transformation of melanocytes/naevi into melanoma (Fan et al., 2011) and was shown to repress the expression of p21/CDKN1A through its interaction with HDAC1. In the context of genotoxic treatment induced senescence in melanoma, the p53 independent expression of p21/CDKN1A could results from the phosphorylation and subsequent degradation of EZH2 by ATM during DDR (Ito et al., 2018). These findings suggest that DNA damaging agents could induce a stable senescence state in melanoma cells, which may contribute to long-term growth arrest and provide potential senolytic intervention windows.

In contrast, long-term treatment with BMi triggered a partially reversible senescence-like phenotype in Braf V600E mutated melanoma cells. While these cells exhibited a G1 growth arrest and increased senescence-associated β-galactosidase activity, they did not display higher levels of DDR foci, secretion of interleukin 8 (IL8) or bcl2-family senolytic sensitivity. We also observed that this induced senescence-like phenotype was partially reversible, as approximately 20% to 30% of the treated cells could regain their proliferative ability and form colonies after drug washout. These observations are consistent with previous reports showing that BRAF and MEK inhibitors induce features of stress-induced senescence, with or without cell death, in BRAF mutant melanoma cells (Haferkamp et al., 2013, Krayem et al., 2018, Li et al., 2016, Madorsky Rowdo et al., 2020, Paraiso et al., 2010, Schick et al., 2015). They observed that BRAF inhibitors, such as vemurafenib and GSK2118436, induce features of stress-induced senescence. In some cases, this phenotype is accompanied by autophagy and characterized by G1 arrest, induction of p27KIP1, and activation of pRb. However, this growth arrest is only transient for some cells that regain their proliferative ability after washing off the BMi. This could be explained, in part, by the rapid recovery of phospho-ERK in some cells, a well-known mechanism for driving therapy escape (Paraiso et al., 2010). Additionally, the autophagy accompanying senescence induced by the inhibitors may reverse senescence in some cells and promote their proliferation after the washout of the inhibitors. Furthermore, prolonged inhibition of BRAF and MEK (8 days or more) leads to a senescence phenotype through Myc degradation and transcriptional upregulation of ERBB3, which is involved in mechanisms of primary resistance to MAPK targeted therapies (Hayes et al., 2016, Sun et al., 2014). We also observed that the longer the BRAF+MEK inhibition was prolonged, the longer it took for cells to reproliferate after drug washout. This suggests that prolonged inhibition could lead to epigenetic changes in cells that remain in a senescence-like state, although that senscent-like state is apparently resistant to bcl2-family senolytics. Intriguingly, a recent study observed that epigenetic inhibitors, such as HDACi and CDK9i, promote the death of senescent melanoma cells surviving long-term BRAF inhibition (Madorsky Rowdo et al., 2020). This supports the idea that long-term BRAF and MEK inhibition can induce a senescence phenotype with unique epigenetic modifications and that specific senolytics targgeting that cell state could be discovered. This distinct phenotype observed in Braf-mutated cells support the idea that the response to therapy depends not only on the cell type but also on the specific treatment modality. These findings highlight the importance of considering the underlying genetic alterations when designing targeted therapies against senescence.

Finally, we investigated the direct cooperation between BCL-2/XL inhibitors and BMi in suppressing melanoma cell viability. Melanoma cells showed sensitivity to Bcl-XL/Bcl-2 inhibitors when exposed simultaneously to BMi, regardless of the cell fate. This synergy has been observed in various contexts, inducing apoptosis and tumor regressions in KRAS mutant cancer models (Corcoran et al., 2013, Koyama et al., 2020). In non-small cell lung cancer, MEK inhibition combined with navitoclax increased apoptotic cells (Tan et al., 2013). MEK and BCL-2/XL inhibitors (GDC-0973 and ABT-263) also inhibited tumor growth in high-grade serous ovarian cancer patient-derived xenograft models (Iavarone et al., 2019). The MAPK pathway, activated in BRAF V600E mutation melanomas, regulates apoptosis through effectors such as BAD and BIM. MEK inhibition enhances Bcl-2/XL inhibitors’ cytotoxicity by promoting BIM dephosphorylation and binding to MCL-1 (Korfi et al., 2016). Current clinical trials are assessing the safety and efficacy of this combination to overcome resistance and relapse in cancer patients (Kun et al., 2021). Our observations confirm the direct synergy between MEK and Bcl-2/XL inhibitors and demonstrate the feasibility of simultaneously inhibition, but reduce the interest of a sequential MEK followed by Bcl-2/XL inhibition in a senescence one-two punch.

In summary, CP and XRA induce senescence in melanoma cells, increasing SASP through DDR and sensitizing the cells to Bcl-2/XL inhibitors. BMi, however, induces a partially reversible senescence-like state non targegtable using current classic senolytics. This partially reversible senescence-like state is consistent with the limited effectiveness of BMi in time leading to high relapse or resistance rates. Further investigations are needed for in vivo validation of melanoma senescence phenotype following various chemotherapies. Nevertheless, sequential combination of senolytics with traditional chemotherapy/radiotherapy or simultaneous combination with BMi holds promise to selectively eliminate senescent cells without harming healthy cells. Senotherapeutics may offer improved outcomes for resistant or secondary melanoma patients. Additionally, exploring the interplay between TIS and the adaptive immune system, (tumor-infiltrating lymphocytes (TILs)) in melanoma or other cancers would be relevant in different therapeutic settings.

## 5 Figures

**Supplementary information 1: Treatments trigger melanoma cell proliferation arrest, morphological changes, and cell death.**

**A)** Table showing IC50 of melanoma cells to different tested drugs. **B)** Immunoblotting illustrating inhibition of MAPK/ERK downstream molecules by BMi 48 hours post-inhibition. **(C-E)** Proliferation curves of melanoma cells exposed to different doses of **C)** Dabrafenib (Braf-i), **D)** Trametinib (Mek-i), and E) the combination of Braf-i+Mek-i. **F)** Flow cytometric analysis of cell cycle distribution 9 days following the indicated treatments in Mel-1102 cells. Mean ± SD of at least 2 independent experiments is shown.

**Supplementary information 2: Therapy-induced senescence phenotypes in melanoma cells include the p21 pathway.**

**A)** Representative images of melanoma cells expressing H2B-GFP (untreated or day 6 following eac-treatment). **B)** Representative images of EdU pulse from day 8 to 9 post-treatment. **(C-F)** Immunoblotting illustrating p53 and p21expression at different time points in **C)** Mel-SK23, **D)** Mel-1102, **E)** Mel 624.38 and **F)** Mel-526 cells exposed to CP or BMi.

**Supplementary information 3: Distinct cells sensitivity to Bcl xl / Bcl-2 inhibitors according to the type of initial treatment**

**(A-D)** Proliferation curves of **A)** Mel-SK23, **B)** Mel-1102, **C)** Mel-624.38 and **D)** Mel-526 cells exposed to a panel of 4 senolytics drugs (ABT-263, A115, ABT-199 and PPL) at different dose range (between 0.16µM to 5µM). **E)** Representative images of cell death (H2BGFP in green; and mask of PI/H2B-GFP double positive in pink) 24 hours following senolytics treatment. ABT-263: BCL-2/XL inhibitor; ABT-199: BCL-2 inhibitor; A-115: BCL-XL inhibitor; PPL: Antioxydant.

## 7 Author Contributions

DT, NM and FR designed the study. DT, JD and NM, performed experiments. DT, NM and FR wrote the manuscript. GC, NM, ST and FR provided technical support, conceptual advice, and revised the manuscript.

## 9 Acknowledgments

We thank former and current members of Dr Turcotte and Dr Rodier Lab for valuable comments and discussions. We thank the Institut du cancer de Montréal (ICM) Imaging and Live imaging platform and the ICM flow cytometry platform.

