## supplemental figs for "Defining melanoma combination therapies that provide senolytic sensitivity in human melanoma cells"

### Supplementary information S1. Treatments trigger melanoma cells growth arrest, morphological changes and cell death

**A**

| | Cell lines | | | |
|---|---|---|---|---|
| | Mel-SK23* | Mel-1102** | Mel-624.38** | Mel-526** |
| **BRAF** | WT | WT | V600E | V600E |
| **NRAS** | WT | Q61K | WT | WT |
| **KRAS** | WT | WT | WT | WT |
| **TP53** | WT | WT | p.Cys275Trp | p.Cys275Trp |
| **Disease site** | Cutaneous | Metastatic | Metastatic | Metastatic |

* Memorial Sloan Kettering Cancer Center; New York; USA
** Rosenberg S.A.; Surgery Branch, National Cancer Institute; NIH; Bethesda; USA

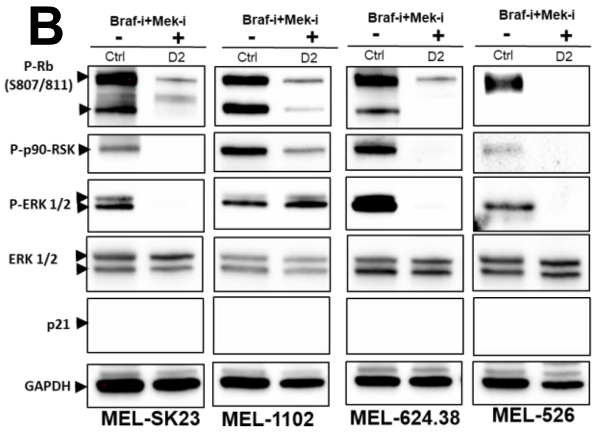


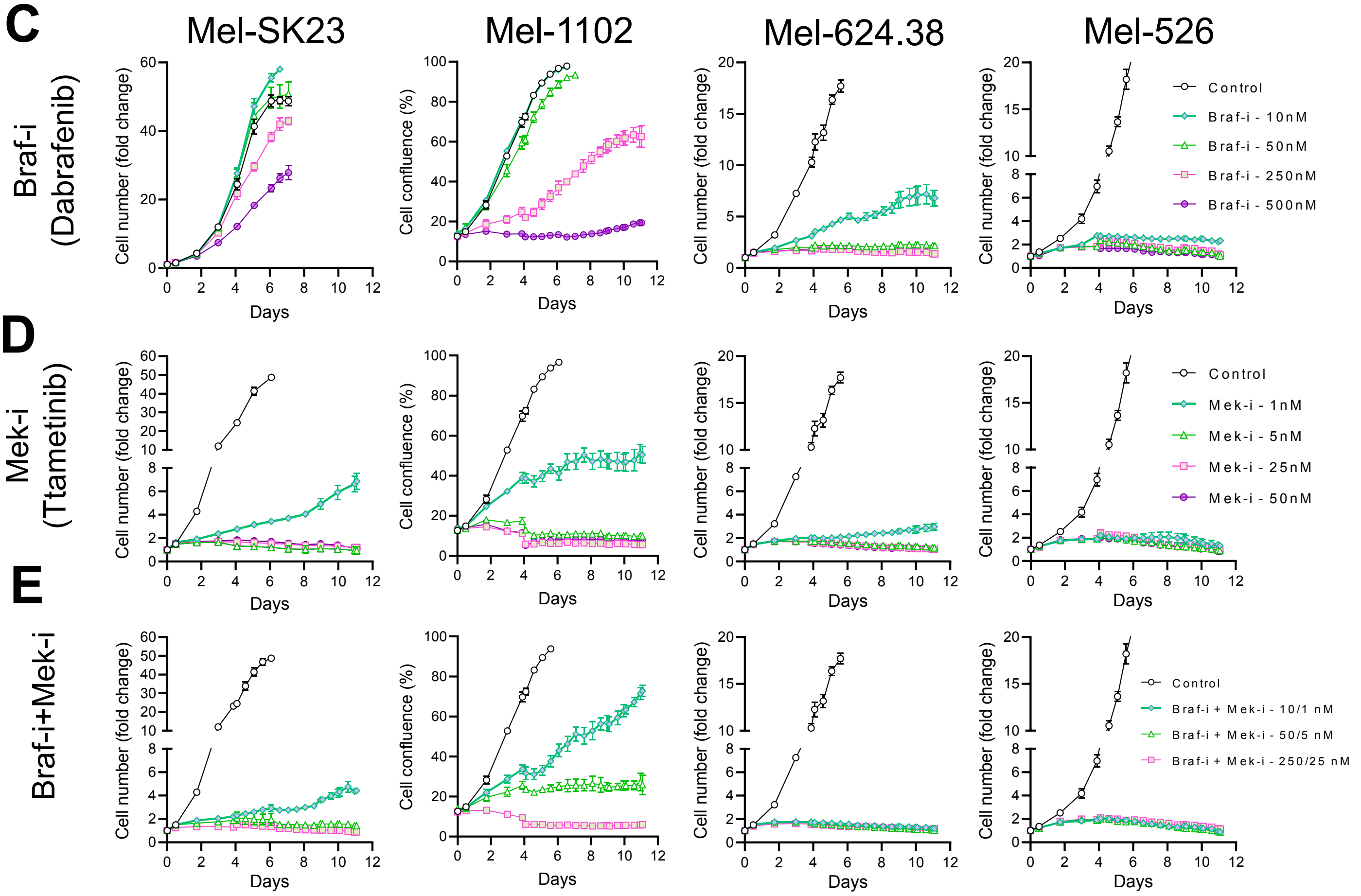


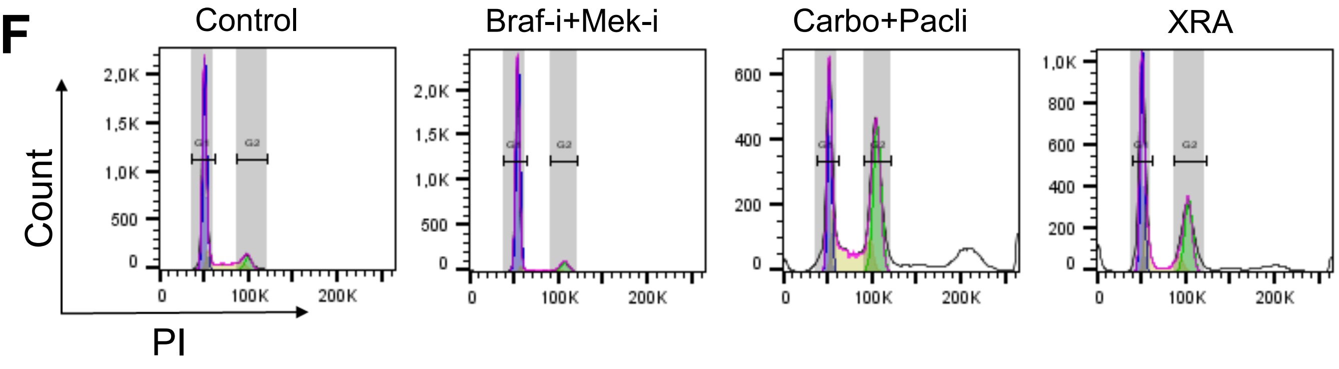

**Supplementary information S2. Therapy induced senescence phenotype in melanoma cells relay on p21 pathway**

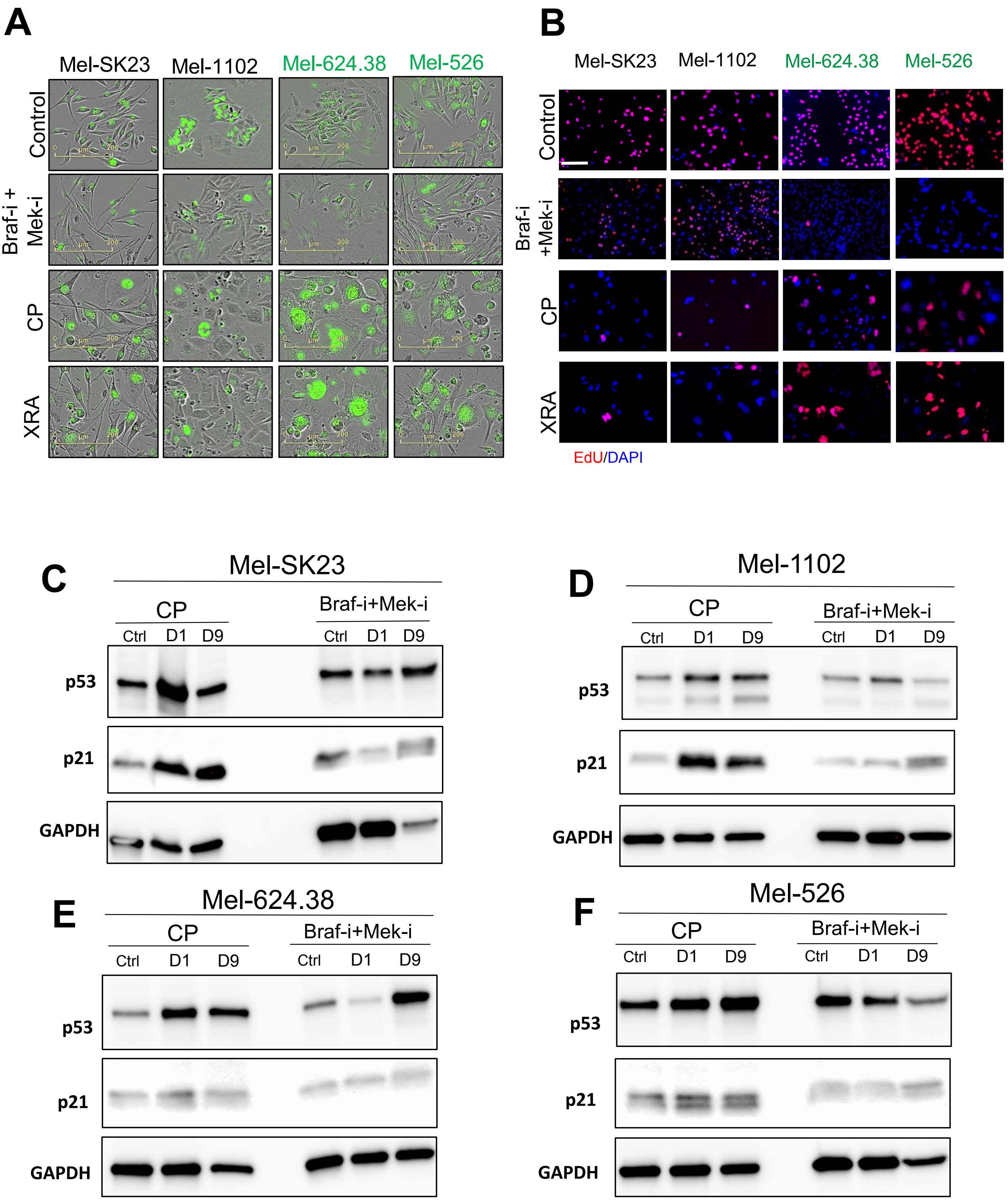

**Supplementary information S3. Distinct cells sensitivity to Bcl xl / Bcl-2 inhibitors according to the type of initial treatment**

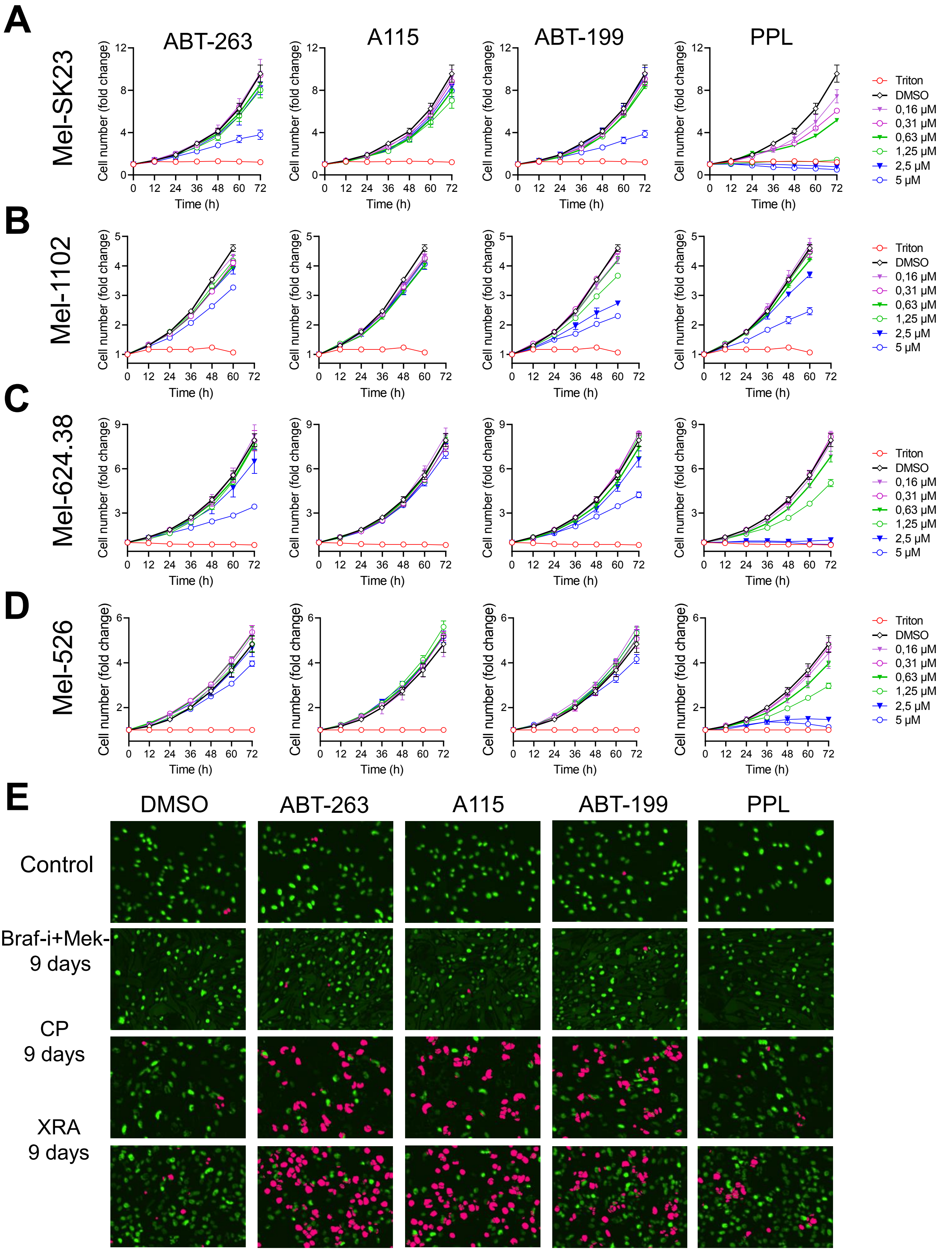
